# Glucocorticoid receptor activation induces gene-specific transcriptional memory and universally reversible changes in chromatin accessibility

**DOI:** 10.1101/2021.01.05.425406

**Authors:** Melissa Bothe, René Buschow, Sebastiaan H. Meijsing

## Abstract

Glucocorticoids are stress hormones that elicit cellular responses by binding to the glucocorticoid receptor (GR), a ligand-activated transcription factor. The exposure of cells to this hormone induces wide-spread changes in the chromatin landscape and gene expression. Previous studies have suggested that some of these changes are reversible whereas others persist even when the hormone is no longer around. However, when we examined chromatin accessibility in human airway epithelial cells after hormone washout, we found that the hormone-induced changes were universally reversed after one day. Reversibility of hormone-induced changes are found for GR-occupied opening sites and also for closing sites that typically lack GR occupancy. These closing sites are enriched near repressed genes, suggesting that transcriptional repression by GR does not require nearby GR binding. Mirroring what we say in terms of chromatin accessibility, we found that transcriptional responses to hormone are universally reversable. Moreover, priming of cells by a previous exposure to hormone, in general, did not alter the transcriptional response to a subsequent encounter of the same cue. Interestingly, despite the short-lived nature of hormone-induced changes in the chromatin landscape, we identified a single gene, *ZBTB16*, that displays transcriptional memory manifesting itself as a more robust transcriptional response upon repeated hormone stimulation. Single-cell analysis revealed that the more robust response is driven by a higher probability of primed cells to activate *ZBTB16* and by a subset of cells that express the gene at levels that are higher than the induction levels observed for naïve cells. Although our study shows that hormone-induced changes are typically reversable, exposure to hormone can induce gene-specific changes in the response to subsequent exposures which may play a role in habituation to stressors and changes in glucocorticoid sensitivity.

## Introduction

Transcriptional memory is an adaptive strategy that allows cells to ‘learn’ from a previous transient exposure to an environmental stimulus and orchestrate a more efficient response when the same cue is encountered again. This phenomenon can manifest as a cell’s ability to elicit a more robust transcriptional response of signal-inducible genes when these cells were primed by a previous encounter with the stimulus (reviewed in [1]). Transcriptional memory has been well-studied in plants that have evolved adaptive transcriptional responses to cope with the various environmental stressors they are subjected to and cannot run away from (reviewed in [2,3]). It has also been described in other systems, such as responses in yeast to environmental signals [4,5] and the response of cells of the immune system to cytokines [6]. The mechanisms that underly these adaptive strategies include altered binding of poised RNA polymerase II as well as changes in the chromatin state, particularly persistent changes in histone modifications, chromatin accessibility and the incorporation of histone variants [1–4,7– 9].

The glucocorticoid receptor (GR) is a hormone-inducible transcription factor (TF) that regulates the expression of genes involved in diverse processes including development, metabolism and immunity (reviewed in [10,11]). Cytoplasmic GR is activated upon the binding of glucocorticoids (GCs), which are secreted from the adrenal cortex in response to various types of stresses including infection, malnutrition and anxiety [12]. Glucocorticoids are also released in a circadian and ultradian manner as short, nearly-hourly pulses (reviewed in [13,14]). Activated GR translocates into the nucleus where it binds to various genomic loci, resulting in the up- or downregulation of its target genes. Extensive research indicates that GR can function as both an activator and repressor of transcription [15]. For transcriptional activation, the paradigm is that direct DNA binding of GR nucleates the assembly of transcription regulatory complexes that modulate target gene expression [15]. For transcriptional repression, the proposed mechanisms responsible are often less clear and more diverse. Some studies suggest that GR downregulates target genes by binding to repressive DNA sequences known as negative glucocorticoid response elements (nGREs) [16,17]. In addition, repression might be mediated by GR tethering to regulatory factors including AP1, NFκB and NGFI-B [18–22]. However, several studies challenge the notion that local occupancy is the general driver of transcriptional repression. Instead, these studies argue that transcriptional repression might be driven by the redistribution of the binding of other transcription factors and coregulators and by alternation of the chromatin structure at enhancers that are not directly occupied by GR [23–26].

Genomic GR binding is associated with a number of changes to the chromatin state [27]. For instance, GR binding induces changes in histone modifications by recruiting cofactors, such as histone methyltransferases and acetyltransferases, that induce post-translational modifications of proteins including histones [15,28–30]. Moreover, GR binding is associated with chromatin remodeling resulting in local increases in chromatin accessibility at its binding sites [31–34]. GR activation also induces chromatin decompaction at a scale that is detectable by light microscopy [35] and stabilizes long-range chromatin interactions as shown by high-throughput sequencing-based methods such as Hi-C [33,36,37].

Interestingly, several studies indicate that GR-induced chromatin changes can persist after the withdrawal of hormone. For example, a genome-wide analysis showed that GR-induced increases in chromatin accessibility are maintained for at least 40 minutes following hormone withdrawal at a subset of GR-binding sites [33]. Another study using a genomically-integrated mouse mammary tumor virus (MMTV)-fragment showed that changes may even persist much longer with GR-induced increases in DNase I-sensitivity of this locus persisting for more than 20 cell divisions after hormone withdrawal [38]. Similarly, large-scale chromatin unfolding that occurred at the *FKBP5* locus upon GR activation persisted for up to 5 days after hormone washout [35], suggesting that GR binding can induce a long-lived chromatin-based ‘memory’ of GR binding. The persistent GR-induced chromatin changes could result in a different transcriptional response of genes when cells are exposed to hormone again. However, if this is indeed the case is largely unknown.

In this study, we sought to (1) link changes in chromatin accessibility upon GC treatment to gene regulation, (2) investigate whether long-lived chromatin changes are a commonly observed feature of GR activation, and (3) whether a previous exposure to GCs can be ‘remembered’ and result in an altered transcriptional response upon a second GC exposure. Our data uncover global increases and decreases in chromatin accessibility that coincide with increased and decreased gene expression, respectively. Even though we find that the changes in chromatin accessibility are universally reversible, we also find indications that cells may ‘remember’ a previous exposure to hormone in a gene-specific manner.

## Methods

### Cell Culture and Hormone Treatments

A549 (ATCC CCL-185) and U2OS-GR cells expressing stably integrated rat GRα [39] were cultured in DMEM supplemented with 5% FBS.

#### Hormone washout treatments in A549 cells (Fig. 2)

Cells were treated with 100 nM dexamethasone (Dex), 100 nM hydrocortisone (Cort) or 0.1% EtOH (as vehicle control) for 20 hours and either harvested immediately or washed 2x with PBS and subsequently cultured in hormone-free medium for 24 hours before harvest.

#### Hormone washout treatments in U2OS-GR cells (Fig. S2)

Cells were treated with 100 nM Dex or 0.1% EtOH for 4 hours and then were either harvested immediately or washed 2x with PBS and subsequently cultured in hormone-free medium for 24 hours before harvest.

#### Hormone re-induction treatments (Fig. 3-6)

Cells were treated with either 100 nM Dex, 100 nM Cort or 0.1% EtOH. After 4 hours, cells were washed 2x with PBS and cultured in hormone-free medium for 24 hours, after which cells were treated again with either 100 nM Dex, 100 nM Cort or 0.1% EtOH.

### RNA extraction and analysis by quantitative real-time PCR (qPCR)

RNA was extracted from cells treated as indicated in the figure legends with the RNeasy Mini kit (QIAGEN) and reverse transcribed using the PrimeScript RT Reagent Kit (Takara). qPCR was performed using primers listed in Table 1.

**Table 1.**
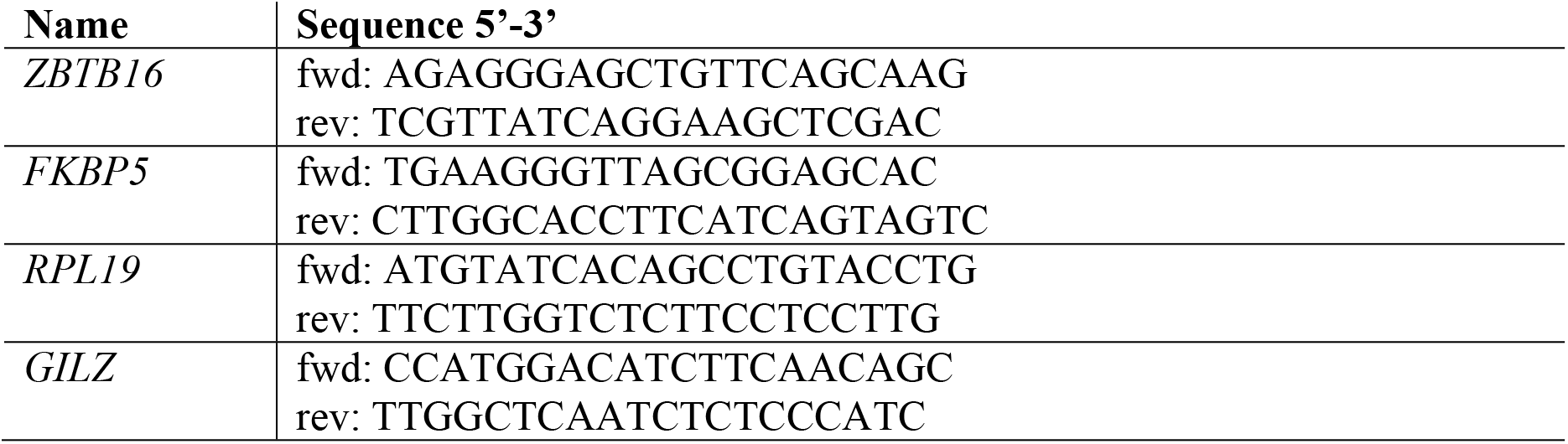
qPCR primers for the quantification of gene expression

### Total RNA-seq

Total RNA was extracted from 1 million A549 cells treated as indicated in the figure legends using the QIAGEN RNeasy Mini kit. Sequencing libraries were prepared with the KAPA RNA HyperPrep kit with RiboErase (Roche #08098131702) and samples were submitted for paired-end Illumina sequencing. RNA-seq data for U2OS-GR cells (1 µM Dex or EtOH, 4h; [40]) were downloaded from ArrayExpress (accession number E-MTAB-6738).

### ATAC-qPCR and ATAC-seq

ATAC assays for A549 and U2OS-GR cells treated as indicated in the figure legends, were performed as previously described [41]. For ATAC-seq, samples were paired-end sequenced. For ATAC-qPCR, ATAC libraries were quantified by qPCR with primers shown in Table 2.

**Table 2:**
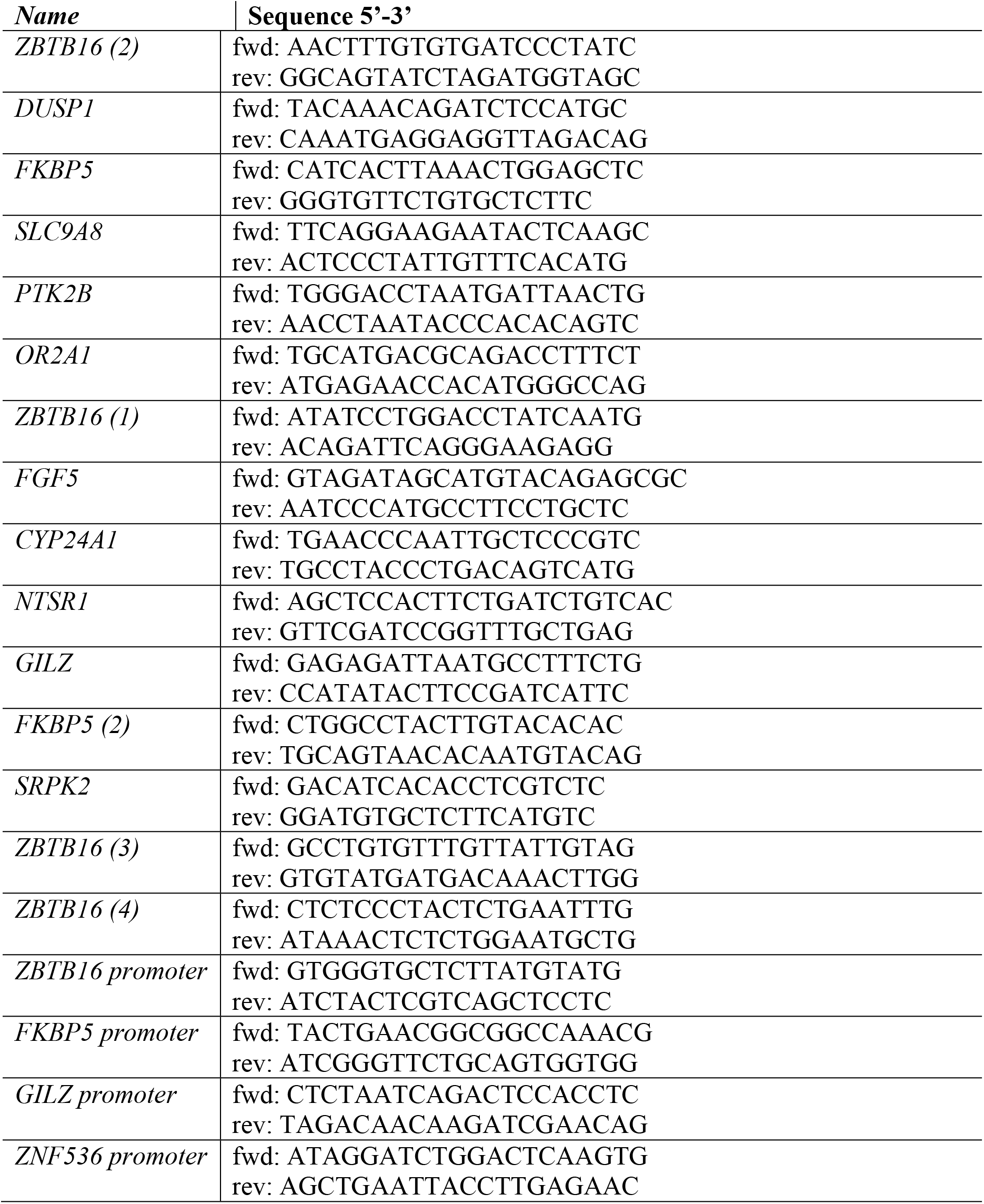
qPCR primers for the quantification of ATAC and ChIP experiments

### ChIP-qPCR and ChIP-seq

ChIP assays for cells treated as indicated in the figure legends, were performed as previously described [42] using antibodies targeting GR (N499, 2 µl/ChIP), H3K27ac (Diagenode #C15410196, 1.4 µg/ChIP), H3K4me3 (Diagenode #C15410003, 1.4 µg/ChIP), H3K27me3 (Diagenode #C15410195, 1.4 µg/ChIP), RNA Polymerase II 8WG16 (Covance #MM2-126R, 2 µg/ChIP) and Phospho-RNA pol II CTD (Ser5) (Thermo Fisher Scientific #MA1-46093, 2 µg/ChIP). ChIP assays for qPCR quantification were performed using primers shown in Table 2. Sequencing libraries were generated with the Kappa HyperPrep kit (Roche #07962363001) and submitted for single-end Illumina sequencing.

For A549 cells, the following ChIP-seq datasets were downloaded from the Gene Expression Omnibus (GEO): GR (2 replicates; 100 nM Dex or EtOH for 3h; GSE79431, [43]), H3K27ac (100 nM Dex, 0h/4h; GSM2421694/GSM2421873; [44]), H3K4me3 (100 nM Dex, 0h/4h; GSM2421504/GSM2421914; [44]), H3K27me3 (100 nM Dex or EtOH, 1h; GSM1003455/GSM1003577; [44]) and p300 (100 nM Dex, 0h/4h; GSM2421805/ GSM2421479; [44]).

For U2OS-GR cells, the following ChIP-seq datasets were downloaded from the Sequence Read Archive (SRA) or ArrayExpress: GR replicate 1 (1 µM Dex, 1.5h; SRA accession SRX256867/SRX256891; [45]), GR replicate 2 (1 µM Dex, 1.5h; ArrayExpress accession E-MTAB-9616; [46], H3K27ac (1 µM Dex or EtOH, 1.5h; ArrayExpress accession E-MTAB-9617; [46]).

### 4C-seq

4C template preparation from 5 million A549 cells treated as indicated in the figure legend was done as described in [47] using Csp6I (Thermo Fisher Scientific) and DpnII (Thermo Fisher Scientific) as primary and secondary restriction enzymes, respectively. Sequencing library preparation was essentially performed as described in [48], with the exception that a 1.5× AMPure XP purification was carried out after the first PCR. Primer pairs used for the inverse PCR are listed in Table 3. 4C libraries were submitted for single-end Illumina sequencing.

**Table 4:**
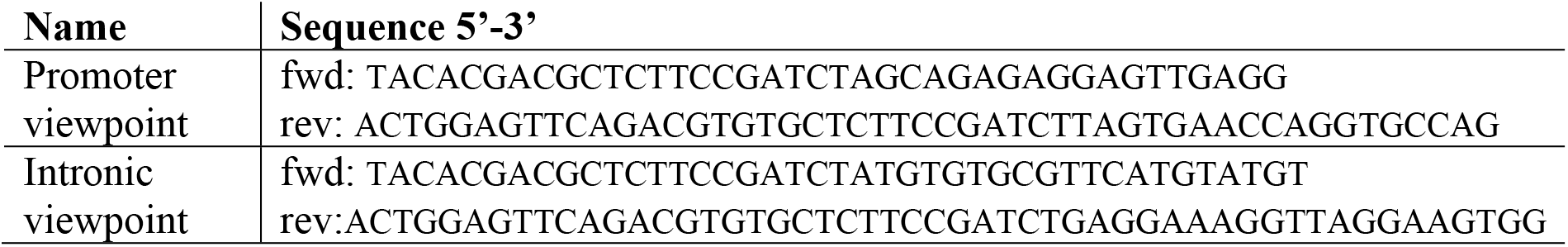
4C primer sequences for inverse PCR

### RNA Fluorescence *in situ* Hybridisation (RNA FISH)

The following FISH probes, labelled with Quasar® 570 Dye, were purchased from Stellaris® (Biosearch Technologies, Inc., Petaluma, CA): (1) human *FKBP5* (#VSMF-2130-5), and (2) human *ZBTB16* which were designed targeting the complete coding sequence of human *ZBTB16* (GenBank: BC029812.1) using the Stellaris® RNA FISH Probe Designer (Biosearch Technologies, Inc., Petaluma, CA) available online at www.biosearchtech.com/stellarisdesigner (version 4.2). The RNA FISH procedure was performed on A549 cells (treated as indicated in the figure legends) according to the manufacturer’s recommendations for adherent cells (www.biosearchtech.com/stellarisprotocols). Images were captured on an Axio Observer.Z1/7 (Zeiss) using a 100x Oil Immersion Objective (NA=1.4) running under ZEN 2.3.

### Computational Analysis

#### ATAC-seq

##### Data processing

Bowtie2 v2.1.0 [49] (--very-sensitive) was used to map the paired-end ATAC-seq reads to the reference human genome hg19. Reads of mapping quality <10 and duplicate reads were filtered out with SAMtools v1.10 [50] and Picard tools (MarkDuplicate) v.2.17.0 (http://broadinstitute.github.io/picard/), respectively. Reads were shifted to account for Tn5 adaptor insertion (described in [51]) with alignmentSieve from deepTools v3.4.1 [52].

##### Calling regions of increasing/decreasing/non-changing accessibility

Regions of increasing/decreasing/non-changing accessibility were determined based on the 20h Dex-, Cort and EtOH-treated (no washout) ATAC-seq data for A549 cells and the 4h Dex- and EtOH-treated (no washout) ATAC-seq data for U2OS-GR cells. For increasing accessibility, peaks were called on hormone-treated samples over vehicle-treated samples with MACS2 v2.1.2 [53] (--broad --broadcutoff 0.001). For sites of decreasing accessibility, peaks were called in vehicle-treated samples over hormone-treated samples using the same MACS2 settings. For A549 cells, called peaks were filtered for a ‘fold_enrichment’ score of >2 and a high-confidence set of increasing/decreasing peaks was obtained by extracting the intersect between Dex- and Cort-treated samples with BEDtools v2.27.1 intersect [54]. For U2OS-GR cells, called peaks were filtered for a ‘fold_enrichment’ score of >3. Sites of increasing (decreasing) accessibility were excluded if they overlapped a promoter region (+/-500 bp around TSS) of any transcript variant of upregulated (downregulated) genes.

To define sites of non-changing accessibility, peaks were called independently for each treatment using MACS2 (--broad --broadcutoff 0.001, ‘no control’). Next, called peaks were filtered for a ‘fold_enrichment’ score of >8 and the intersect between all treatments was extracted and sites of increasing and decreasing accessibility were removed with BEDtools intersect. ENCODE blacklisted regions for hg19 [44] and regions within unplaced contigs and mitochondrial genes were removed from all peaks.

To define sites of increasing/decreasing accessibility before and after washout, regions of increased or decreased accessibility were called using the ‘24h after washout’ ATAC-seq data sets in the same way as described above for the ‘before washout’ samples. Subsequently, overlapping sites between the ‘before’ and ‘after washout’ were extracted with BEDtools v2.27.1 intersect.

##### Normalization for heatmap and genome browser visualizations

To account for differences in signal-to-noise ratios, ATAC-seq samples were normalized with individual scaling factors. For this purpose, peaks were called for each treatment using MACS2 v2.1.2 (--broad --broadcutoff 0.01, ‘no control’). To obtain a list of ATAC-seq peaks of high-confidence, the intersect of all treatments was extracted with BEDtools intersect. Having removed regions within unplaced contigs and mitochondrial DNA as well as ENCODE blacklisted regions for hg19, fragments of each sample were counted on the high-confidence regions using featureCounts (allowMultiOverlap=TRUE, isPairedEnd=TRUE) [55]. The estimateSizeFactorsForMatrix function from DEseq2 v1.26.0 [56] was applied to calculate the scaling factors. The reciprocals of the resulting factors were taken and provided as scaling factors to the deepTools v3.4.1 [52] function bamCoverage to generate scaled bigWigs. Heatmaps and mean signal plots (+/-2 kb around the peak center) were generated with the deepTools functions computeMatrix (reference-point) and plotHeatmap, using the scaled bigWig files as input.

#### ChIP-seq

Bowtie2 v2.1.0 [49] (--very-sensitive) was used to map ChIP-seq reads to the reference genome hg19. The GR, ChIP-seq reads for rep1 (SRP020242, [45]) were mapped by setting options in Bowtie2 as ‘--very-sensitive -X 600 --trim5 5’. SAMtools v1.10 [50] was utilized to remove reads of mapping quality <10. Duplicate reads were filtered out using the MarkDuplicate function from Picard tools v.2.17.0 (http://broadinstitute.github.io/picard/). RPKM-normalized bigWig files were generated with bamCoverage from deepTools v3.4.1 [52]. Heatmaps and mean signal plots (+/-2 kb around the peak center) were generated with the functions computeMatrix (reference-point) and plotHeatmap from deepTools. GR ChIP-seq peaks for each replicate were called over input with MACS2 v2.1.2 [53] setting a qvalue cut-off of 0.01. BEDtools intersect v2.27.1 (-u) [54] was used to extract overlapping peaks that were called in both replicates to obtain a final GR peak set. ENCODE blacklisted regions for hg19 [44] and regions within unplaced contigs and mitochondrial genes were removed.

#### RNA-seq

Paired-end reads were aligned to the hg19 reference genome using STAR v2.7.0a [57]. Reads of mapping quality <10 were removed with SAMtools v1.10 [50]. For genome browser visualization, triplicates were merged using the merge function from SAMtools and RPKM-normalized bigWig files were generated using bamCoverage from deepTools v3.4.1 [52].

Differential gene expression analysis for A549 cells was performed based on the quantification of read coverage in introns of the total RNA-seq data. For this purpose, the annotation file of NCBI RefSeq genes available from the UCSC Genome Browser [58] was downloaded and information on the longest transcript variants per gene were extracted. Introns of the longest transcripts were obtained with the intronicParts function from GenomicFeatures [59]. Next, to ensure that the intronic regions do not overlap any mRNA sequences, exonic regions of all transcripts (obtained with the exonicParts function from GenomicFeatures) were subtracted from the introns (using the GenomicFeatures function disjoin on the combined intronic and exonic regions followed by subsetByOverlaps). Finally, introns which associated with more than one gene were excluded to ensure only unique intron regions were contained in the final set. Reads within intronic regions were counted with featureCounts [55] (isPairedEnd=TRUE, primaryOnly=TRUE, requireBothEndsMapped=TRUE, countChimericFragments=FALSE, useMetaFeatures=FALSE). Next, the intronic read counts per gene were summed. Differential expression analysis was performed using DESeq2 v1.26.0 [56]. For U2OS-GR cells, mRNA-seq data was from ArrayExpress accession number E-MTAB-6738 and differential expression analysis was carried out based on exonic read coverage. Exonic regions of the longest transcripts were obtained with the exonicParts(linked.to.single.gene.only=TRUE) and disjoin functions from GenomicFeatures [59]. Read counting and differential expression analysis were performed as described above for the A549 cells.

### Linking peaks to gene regulation

Genes were grouped as upregulated (log2 fold change > 1, adjusted p-value < 0.05, baseMean > 40), downregulated (log2 fold change < −1, adjusted p-value < 0.05, baseMean > 40) or non-regulated (0.1 > log2 fold change > −0.1, baseMean > 40). For nonregulated genes, 500 genes were randomly sampled for A549 cells and 1000 genes for U2OS-GR cells. For each gene, it was determined whether at least one peak fell within a +/-50 kb window around the TSS of the longest transcript variant using BEDtools intersect v2.27.1 (-u) [54]. If a peak overlapped the +/-50 kb window around the TSS of multiple genes it was assigned to the closest gene. P-values were calculated performing a Fisher’s exact test.

#### 4C-seq

4C-seq data were analyzed with the pipe4C pipeline [48] using default settings except setting the --wig parameter to obtain WIG output files.

### RNA FISH Image Analysis

Raw images were processed by applying a Maximum Intensity Projection (MIP) in ZEN 3.0 (Zeiss) including the full Z-Stack of 26 slices. To count transcription sites and individual transcripts, image analysis was performed in ZEN 3.0. Specifically, nuclei were detected by the DAPI staining using fixed intensity thresholds after a faint smoothing (Gauss: 2.0) and segmentation (watershed: 10), and subsequently filtering by circularity (0.5-1), size (75-450 µm^2^) and a mean intensity of maximum (4200). To define a region for the cytoplasm, a ring (width 30 pix = 3.96 µm) was automatically drawn around the nuclei. Transcripts were identified within the nuclei and surrounding rings after Rolling Ball Background Subtraction with a radius 5 pix by a fixed fluorescence threshold and subsequently filtered by area (0-0.33 µm^2^). Transcription sites were identified with the same parameters as the transcripts as well as additionally filtering for an area larger than 0.38 µm^2^.

## Data availability

The ATAC-seq, RNA-seq, ChIP-seq and 4C-seq data generated for this study were submitted to the ArrayExpress repository under the following accession numbers:

Experiment ArrayExpress accession: E-MTAB-9911

Title: Studying changes in chromatin accessibility in A549 cells by ATAC-seq directly after 4-hour glucocorticoid treatment and following a 24-hour hormone washout

Username: Reviewer_E-MTAB-9911 Password: dUyfiojr

Experiment ArrayExpress accession: E-MTAB-9910

Title: Studying changes in chromatin accessibility in A549 cells by ATAC-seq directly after 20-hour glucocorticoid treatment and following a 24-hour hormone washout

Username: Reviewer_E-MTAB-9910 Password: zuE742zz

Experiment ArrayExpress accession: E-MTAB-9909

Title: Studying changes in chromatin accessibility in U2OS-GR cells by ATAC-seq directly after 4-hour glucocorticoid treatment and following a 24-hour hormone washout

Username: Reviewer_E-MTAB-9909 Password: oXRLH12W

Experiment ArrayExpress accession: E-MTAB-9912

Title: Studying changes in chromatin accessibility in A549 cells by ATAC-seq after glucocorticoid re-induction

Username: Reviewer_E-MTAB-9912 Password: cfnyym3p

Experiment ArrayExpress accession: E-MTAB-9914

Title: Glucocorticoid receptor profiling (ChIP-seq) in U2OS-GR cells 24 hours after hormone washout

Username: Reviewer_E-MTAB-9914 Password: faovxysz

Experiment ArrayExpress accession: E-MTAB-9915

Title: Circularized chromosome conformation capture (4C-seq) in A549 cells

Username: Reviewer_E-MTAB-9915

Password: 78u1MzpV

Experiment ArrayExpress accession: E-MTAB-9923

Title: Total RNA-seq in the A549 cell line after glucocorticoid re-induction

Username: Reviewer_E-MTAB-9923

Password: yKKHB56c

## Results

### Linking GR occupancy, chromatin accessibility and gene regulation

To better understand the link between GR binding and changes in chromatin accessibility, we mapped genome-wide changes in chromatin accessibility upon GC treatment in A549 cells by ATAC-seq. Specifically, we analyzed changes that occur following a 4-hour or a 20-hour treatment with either dexamethasone (Dex), a synthetic GC, or with the natural hormone hydrocortisone (Cort). In agreement with previous studies [31–34], many sites showed an increase in chromatin accessibility (‘opening sites’ for both hormone treatments when compared to the control treatment (Fig. 1a,b, S1a)). Interestingly, chromatin accessibility was reduced for an even larger number of sites (‘Closing sites’, Fig. 1a,b, S1a), corroborating findings in macrophages that also reported both increases and decreases in chromatin accessibility upon GC treatment [23]. We could validate several examples of opening and closing sites and noticed that opening sites are often GR-occupied whereas closing sites are not occupied by GR (Fig. 1b,c). For a systematic analysis of the link between GR occupancy and chromatin accessibility changes, we integrated the ATAC-seq results with available GR ChIP-seq data [43]. This analysis showed that GR binding is observed at the majority of opening sites (Fig. 1d). In contrast, only a minor subset of closing sites shows GR ChIP-seq signal (Fig. 1d). Similarly, H3K27ac levels, a mark of active chromatin, increase at opening sites whereas levels decrease at closing sites upon hormone treatment (Fig. 1e). In addition, H3K27ac levels show a modest decrease at sites of non-changing accessibility, suggesting that GR activation induces a global redistribution of H3K27ac (Fig. 1e). These findings are indicative of a re-distribution of the enzymes (*e*.*g*. p300, [60]) that deposit the H3K27ac mark upon GC treatment, which has been described in a mouse mammary epithelial cells [61] and for the estrogen receptor, a GR paralog [62]. To test the effect of GC treatment on global p300 binding in A549 cells, we intersected our data with published p300 ChIP-seq data [44]. This analysis showed a marked increase in p300 signal at GR-occupied opening sites whereas p300 is lost from closing sites that are typically not GR occupied (Fig. S1b).

**Figure 1.**
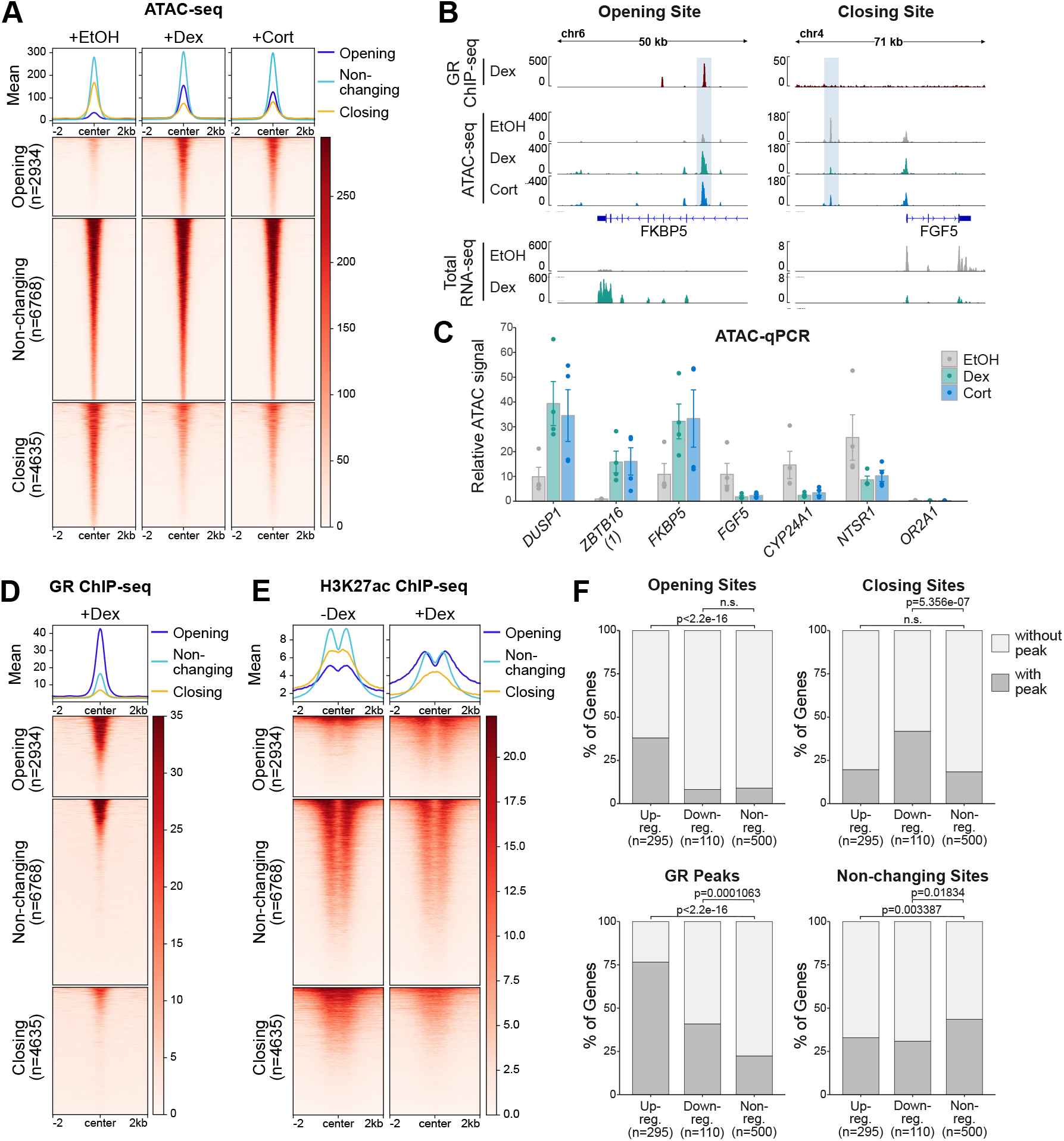
Integrated analysis of genome-wide changes in chromatin accessibility and transcript levels upon GR activation. (a) Heatmap visualization and mean signal plot of normalized ATAC-seq read coverage in A549 cells at sites of increasing (‘opening’), non-changing (‘non-changing’) and decreasing (‘closing’) chromatin accessibility upon hormone treatment (+/-2 kb around center). Cells were treated with EtOH vehicle (20h), Dex (100 nM, 20h) or Cort (100 nM, 20h). (b) Genome browser visualizations of the *FKBP5* and *FGF5* loci in A549 cells showing GR ChIP-seq (100 nM Dex, 3h; RPKM-normalized; data from [43]), ATAC-seq (normalized) and RNA-seq (RPKM-normalized, merge of three replicates) signal tracks. For ATAC-seq experiments, cells were treated with EtOH (20h), Dex (100 nM, 20h) or Cort (100 nM, 20h). For RNA-seq, cells were treated as detailed in Fig. 3a, receiving a vehicle control treatment (4h), followed by a 24h-hormone-free period and a subsequent 4h Dex-(100 nM) or EtOH-treatment (‘--’: EtOH, ‘-+’: Dex). Opening/closing site is highlighted with blue shading. ATAC-qPCR at sites opening or closing upon hormone treatment near indicated genes in A549 cells. Cells were treated with EtOH (20h), Dex (100 nM, 20h) or Cort (100 nM, 20h). Mean ATAC signal (normalized to gDNA) ± SEM (n = 4) is shown. (d) Same as for (a) except that RPKM-normalized GR ChIP-seq read coverage (100 nM Dex, 3h) in A549 cells is shown (data from [43]). (e) Same as for (a) except that RPKM-normalized H3K27ac ChIP-seq read coverage (+/-100 nM Dex, 4h) in A549 cells is shown (data from [44]). (f) Stacked bar graphs showing the percentage of genes in A549 cells of each category (upregulated, downregulated and nonregulated) that have at least one peak for each type (opening, closing and non-changing sites and GR peaks) within +/-50 kb around the TSS. P-values were calculated using a Fisher’s exact test. n.s.: not significant.

**Figure 2.**
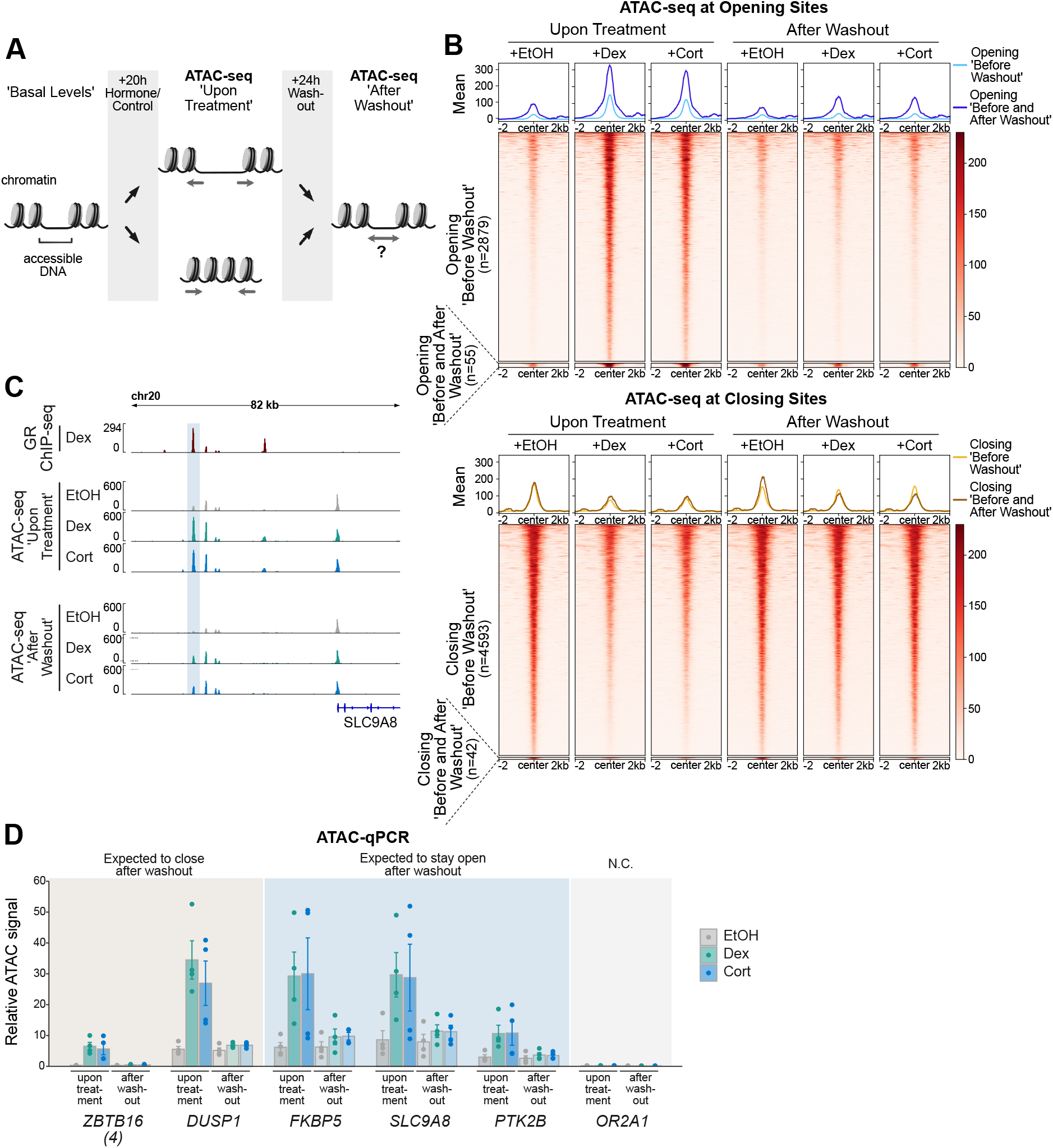
GC-induced changes in chromatin accessibility are universally reversable. (a) Cartoon depiction of the experimental design to assess changes in chromatin accessibility upon hormone treatment and following hormone washout. ATAC-seq was performed (1) on A549 cells treated with Dex/Cort (100 nM) or EtOH for 20h (‘upon hormone treatment’) and (2) on cells treated with Dex/Cort (100 nM) or EtOH for 20h, followed by hormone washout and incubation in hormone-free medium for 24h (‘after washout’). (b) Heatmap visualization and mean signal plot of normalized ATAC-seq read coverage at opening sites (top) and closing sites (bottom) (+/-2 kb around center). Cells were treated as described in (a). The regions are divided into sites which show reversible increased/decreased accessibility upon hormone treatment and regions with persistent changes (opening ‘Before and After Washout’). (c) Genome browser visualization of the *SLC9A8* locus in A549 cells showing GR ChIP-seq (100 nM Dex, 3h; RPKM normalized; data from [43]) and ATAC-seq (normalized) signal tracks. For ATAC-seq, cells were treated as described in (a). Candidate site with persistent increased accessibility after hormone washout is highlighted. (d) ATAC-qPCR of sites opening upon hormone treatment near indicated genes in A549 cells. Cells were treated as described in (a). Regions which are expected (based on the ATAC-seq data) to close or remain accessible following hormone washout are indicated. Mean ATAC signal (normalized to gDNA) ± SEM (n = 4) is shown. N.C. Negative Control.

**Figure 3.**
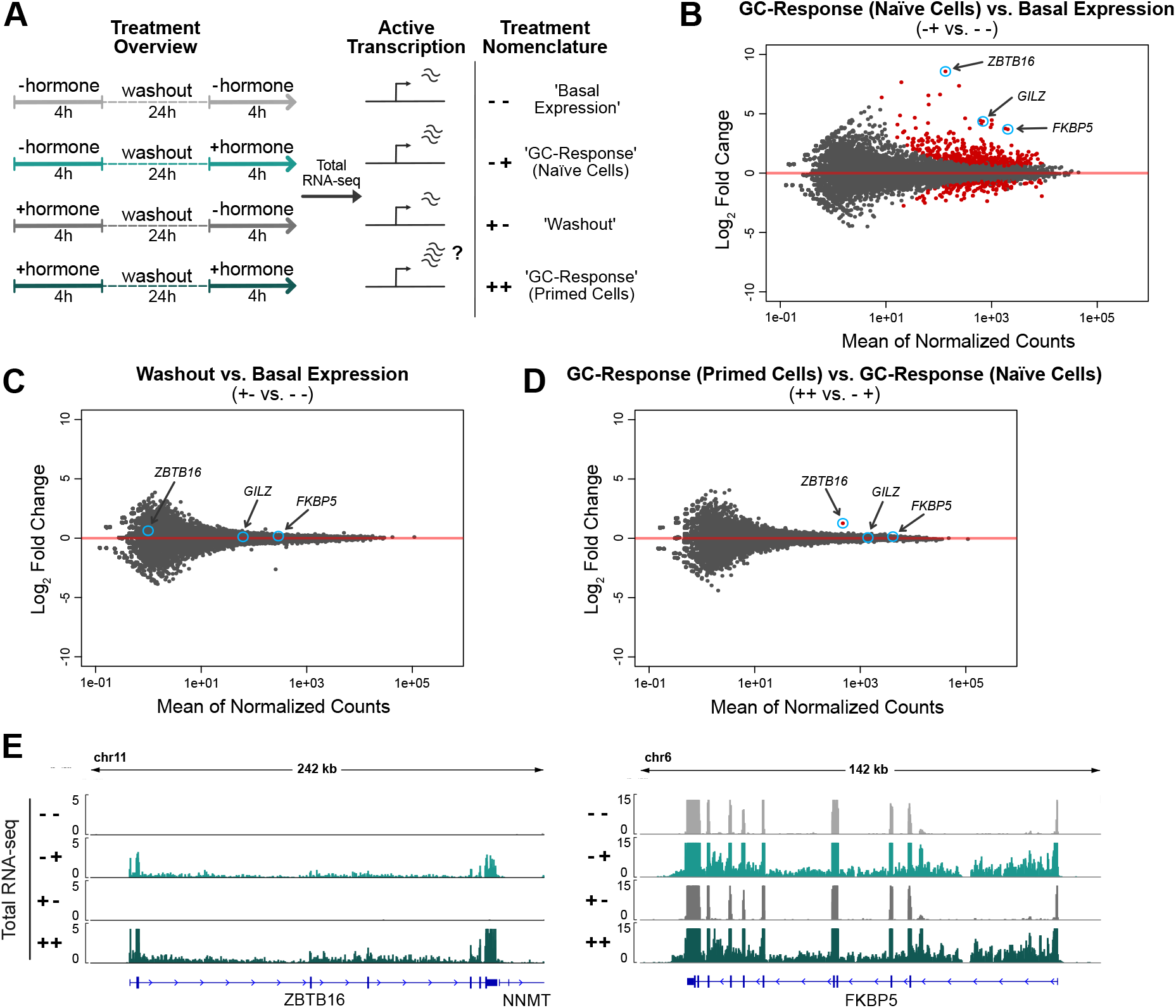
Priming results in a more robust transcriptional response of the *ZBTB16* gene upon repeated GC exposure. (a) Cartoon illustrating our experimental set-up to study how priming by a previous hormone treatment influences a subsequent transcriptional response to hormone. Cells were initially treated with 100 nM Dex (‘primed’) or EtOH (naïve) for 4h. Subsequently, hormone was washed out and cells were cultured in hormone-free medium for 24h, after which cells were treated again with either Dex (100 nM) or EtOH for 4h. (b) MA-plots illustrating the mean of normalized counts and the log2 fold changes for genes when comparing cells subjected to the following treatments: ‘-+’ vs. ‘--’(c) ‘+ -’ vs. ‘--’ (d) ‘++’ vs. ‘-+’. Differential expression analysis was carried using total RNA-seq data and quantifying read coverage within introns. The cells were treated as described in (a). Red dots represent genes significantly up- or downregulated (FDR < 0.001). (e) Genome browser visualizations of the *ZBTB16* and *FKBP5* loci in A549 cells showing total RNA-seq (RPKM-normalized, merge of three replicates) signal tracks. The cells were treated as described in (a).

Next, we set out to assess the link between changes in chromatin accessibility, GR binding and gene regulation. Therefore, we generated total RNA-seq data and analyzed read coverage at introns as a proxy for nascent transcript to capture acute transcriptional responses [63] and defined three categories of genes: upregulated (295), downregulation (110) and nonregulated (randomly sampled 500). For each gene category, we scanned a window of 100 kb centered on the transcription start site (TSS) of each gene for opening, closing and non-changing ATAC-seq peaks while removing sites of increasing accessibility that overlapped TSSs of upregulated genes and sites of decreasing accessibility that overlapped TSSs of downregulated genes. Consistent with expectation, opening peaks and GR peaks are enriched near upregulated genes (Fig. 1f). Conversely, downregulated genes are enriched for closing peaks. However, in contrast to upregulated genes, downregulated genes only show a modest enrichment of GR peaks. Similarly, analysis of a U2OS cell line stably expressing GR (U2OS-GR, [39]) showed only a modest enrichment of GR peaks near downregulated genes (Fig. S1f). Furthermore, closing peaks, which show GC-induced loss of H3K27ac levels and lack GR occupancy (Fig. S1c-f), were enriched near repressed genes.

Taken together, our results further support a model put forward by others [23,24,61,64] in which transcriptional activation by GR is driven by local occupancy whereas transcriptional repression, in general, does not require nearby GR binding. Instead, our data support a ‘squelching model’ whereby repression is driven by a redistribution of cofactors away from enhancers near repressed genes that become less accessible upon GC treatment yet lack GR occupancy.

### GR-induced changes in chromatin accessibility are universally reversible

A previous study reported that a subset of opening sites remains open 40 minutes after hormone withdrawal, indicating that cells retain a ‘memory’ of previous hormone exposure [33]. Memory may even be ‘long-term’ given that a locus-specific study with a genomically-integrated mouse mammary tumor virus showed GC-induced opening that persisted for more than 9 days [38]. Here, we set out to test if long-term memory of opening sites is a general phenomenon and to expand the analysis to loci with reduced chromatin accessibility upon GC treatment. To mirror the study that reported persistent changes after more than 9 days [38], we treated A549 cells for 20 hours with either Dex, Cort or EtOH as vehicle control followed by a hormone-free washout period (Fig. 2a). However, instead of 9 days we decided to assay chromatin accessibility by ATAC-seq after a 24-hour washout period which we reasoned would be more likely to reveal persistent changes. Analysis of the ATAC-seq data showed that the vast majority of opening sites revert to their untreated chromatin accessibility levels (2879/2934 opening sites, Fig. 2b). Similarly, the vast majority of closing sites are transiently closed upon hormone exposure (4593/4635 closing sites, Fig. 2b). To test if the small number of opening sites with apparent persistent increases in chromatin accessibility were true or false positives of our genome-wide analysis, we performed ATAC followed by qPCR on sites which exhibited the most pronounced residual accessibility (example of a candidate site with persistent opening shown in Fig. 2c). The ATAC-qPCR results, however, did not validate the existence of maintained accessibility for these sites and any differences in ATAC signal between basal levels and following hormone washout were, if present at all, subtle (Fig. 2d). To test if persistent GC-induced changes in chromatin accessibility can be observed in another cell type, we performed ATAC-seq after Dex or EtOH and subsequent washout in U2OS-GR cells. Consistent with our findings in A549 cells, we found that the vast majority of both opening and closing sites revert to their initial state after hormone withdrawal (Fig. S2a). However, in contrast to our findings in A549 cells, we could validate maintained accessibility at candidate loci by ATAC-qPCR in U2OS-GR cells (Fig. S2c). In addition, H3K27ac ChIP experiments showed that GC-induced increases in H3K27ac levels were maintained 24 hours after hormone withdrawal at persistent opening sites whereas they reversed to their basal level for sites with transient opening (Fig. S2c). To determine whether GR still occupies persistent opening sites, we performed GR ChIP and ChIP-seq on cells 24 hours after hormone withdrawal. As expected, we found that GR occupancy was lost at sites with transient opening (Fig. S2c,d). In contrast, GR still occupied persistent opening sites 24 hours after hormone withdrawal (Fig. S2c,d) indicating that the sustained accessibility is likely the result of residual GR binding to those sites. One explanation for the sustained occupancy at persistent loci is that a small fraction of GR is still hormone-occupied despite the 24-hour washout and dissociation half-life of Dex of approximate 10 minutes [65]. Interestingly, at low (sub-*K*_*d*_), Dex concentrations, GR selectively occupies a small subset of the genomic loci that are bound when hormone is present at saturating concentrations [66]. To test if GR preferably binds at persistent sites at low hormone concentrations, we performed GR ChIPs at different Dex doses (Fig. S2e) and found residual occupancy at persistent opening sites at very low Dex concentrations (0.1 nM), whereas GR occupancy was no longer observed at sites with transient opening (Fig. S2e). This suggests that the few sites with persistent opening are likely a simple consequence of an incomplete hormone washout and associated residual GR binding.

Together, we conclude that GC-induced changes in chromatin accessibility are universally reversible upon hormone withdrawal with scarce signs of ‘long-term’ memory of previous hormone exposure.

### Prior exposure to GCs results in a more robust regulation of the GR-target gene *ZBTB16*

Next, we were interested to determine if a previous exposure to GCs influences the transcriptional response to a second exposure of the same cue. Therefore, we compared the transcriptional response to a 4-hour Dex treatment between ‘naïve’ cells and cells that were ‘primed’ by a prior 4-hour Dex treatment followed by a 24-hour-hormone-free recovery phase (Fig. 3a). To capture acute transcriptional responses, we performed total RNA-seq and analyzed read coverage at introns to capture nascent transcripts [63]. As expected, we found that GC treatment of naïve cells resulted in the up- or downregulation of many transcripts (Fig. 3b). Importantly, comparison of the basal expression levels between primed cells and naïve cells showed that nascent transcript levels did not show significant changes for any gene indicating that the transcriptional changes were universally reversed after the 24-hour-hormone-free washout (Fig. 3c). When comparing the transcript levels for Dex-treated cells between naïve and primed cells, we found that transcript levels were essentially the same with a single significant exception, the GR-target gene *ZBTB16* (zinc finger and BTB domain containing gene 16, Fig. 3d,e). To further substantiate this finding, we performed qPCR which confirmed that a prior hormone treatment resulted in a more robust upregulation of *ZBTB16* mRNA levels whereas prior treatment did not change the upregulation of two other GR target genes, *GILZ* and *FKBP5* (Fig. 4). Notably, increased mRNA levels are not a simple consequence of mRNA accumulation given that mRNA levels after washout (+-treatment condition) return to the levels observed for untreated naïve cells (Fig. 3e, Fig. 4a). A more robust *ZBTB16* activation upon GC treatment was also observed when cells were primed followed by a longer, 48-hour-hormone-free recovery phase (Fig. S3a), suggesting that cells ‘remember’ a prior hormone exposure through a cell division cycle. Moreover, *ZBTB16* upregulation was further enhanced when cells were exposed to GCs a third time (Fig. 4b). To test if priming of the *ZBTB16* gene is cell type-specific, we analyzed how priming influences a subsequent response to hormone in U2OS-GR cells. When we used Dex to prime U2OS-GR cells, we observed that basal *ZBTB16* levels remained high after hormone washout consistent with residual GR occupancy (Fig. S3c). Therefore, we now used Cort, which has a shorter dissociation half-life than Dex [65], to prime cells. Consistent with our observations using Dex, we found that priming A549 cells with Cort resulted in a more robust upregulation of the *ZBTB16* gene upon a subsequent hormone exposure (Fig. S3b). In contrast, priming of U2OS-GR cells did not alter the response of *ZBTB16* to a subsequent hormone exposure (Fig. S3d).

**Figure 4.**
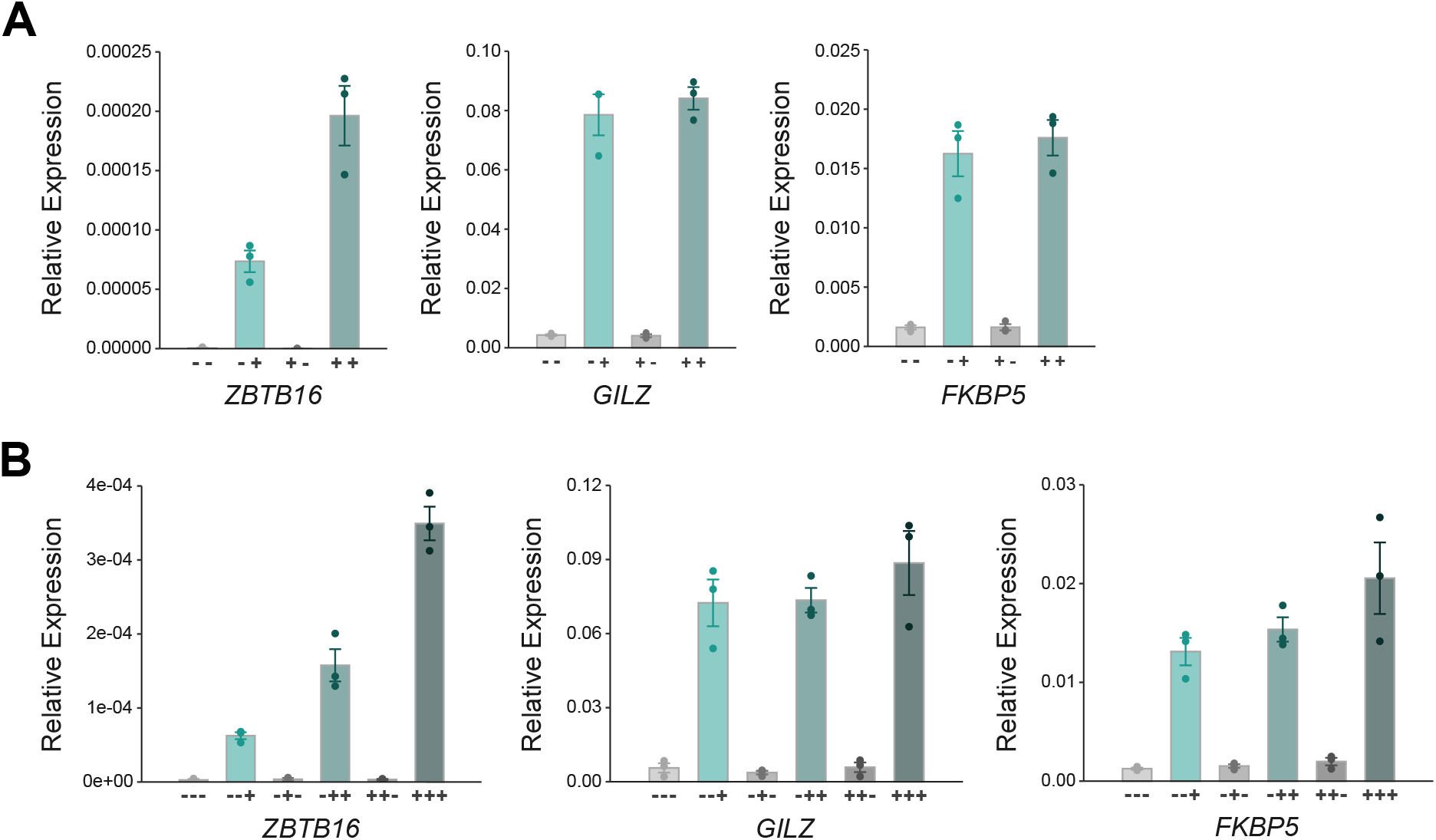
Validation that priming results in a more robust transcriptional response of the *ZBTB16* gene. (a) RT-qPCR results for GR-target genes in A549 cells. The cells were treated as detailed in (Fig. 3a). Mean expression relative to *RPL19* ± SEM (n = 3) is shown. (b) RT-qPCR results for *ZBTB16* in A549 cells. Cells were treated similarly as detailed in (Fig. 3a), except that they received three rounds of hormone treatment: after the second treatment, cells were subjected to washes and cultured in hormone-free medium, and treated again 48h after the first washout with either Dex (100 nM) or EtOH for 4h. Mean relative expression to *RPL19* ± SEM (n = 3) is shown.

Together, our results indicate that a previous exposure to GCs does not result in large-scale reprogramming of the transcriptional responses to a second exposure, yet argue for cell type-specific transcriptional memory for the *ZBTB16* gene.

### Short-term maintenance of GC-induced changes in chromatin accessibility, 3D genome organization and PolII occupancy

To determine if persistent changes in the chromatin state contribute to transcriptional memory of the *ZBTB16* gene, we investigated changes in (1) H3K4me3 and H3K27me3 levels, (2) chromatin accessibility, (3) RNA polymerase II (PolII) occupancy at the promoter, and (4) 3D chromatin organization. Analysis of available ChIP-seq data for A549 cells [44] showed that *ZBTB16* is situated within a repressed genomic region, with high H3K27me3 levels and low or absent H3K4me3 and H3K27ac signal, respectively (Fig. 5a). However, ChIP-experiments we performed showed that priming did not result in significant changes in H3K4me3 or H3K27me3 levels for either treated cells or after hormone washout indicating that these marks likely do not contribute to the transcriptional memory observed (Fig. 5b). Similarly, increases in chromatin accessibility at GR-occupied loci that occur near the *ZBTB16* gene reverted to their uninduced levels after hormone washout (Fig. 5d,e). Furthermore, although priming resulted in the accumulation of PolII and PolII phosphorylated at serine 5 at the promoter of *ZBTB16*, we did not detect any perceptible maintenance of this accumulation after washout (Fig. 5c). Next, we decided to investigate a possible role of 3D chromatin organization, given that GR activation results in chromatin decompaction that persisted for 5 days [35]. Moreover, a recent study into the mechanisms responsible for priming by interferons indicated a role for cohesin and topologically associating domain (TADs) in transcriptional memory [6]. To probe for changes in 3D organization, we performed circular chromosome conformation capture (4C) experiments [47] using either the promoter of the *ZBTB16* gene or an intronic GR-occupied region as viewpoint. The 4C-seq experiments showed an increase in the relative contact frequency between the *ZBTB16* promoter and a cluster of intronic GR binding sites upon hormone treatment (Fig. 5f). However, after washout this increased contact frequency was reversed and furthermore showed comparable levels for primed cells (Fig. 5f), indicating that changes in long-range chromatin interactions at the *ZBTB16* locus upon hormone induction are not maintained and likely do not explain the transcriptional memory observed.

**Figure 5.**
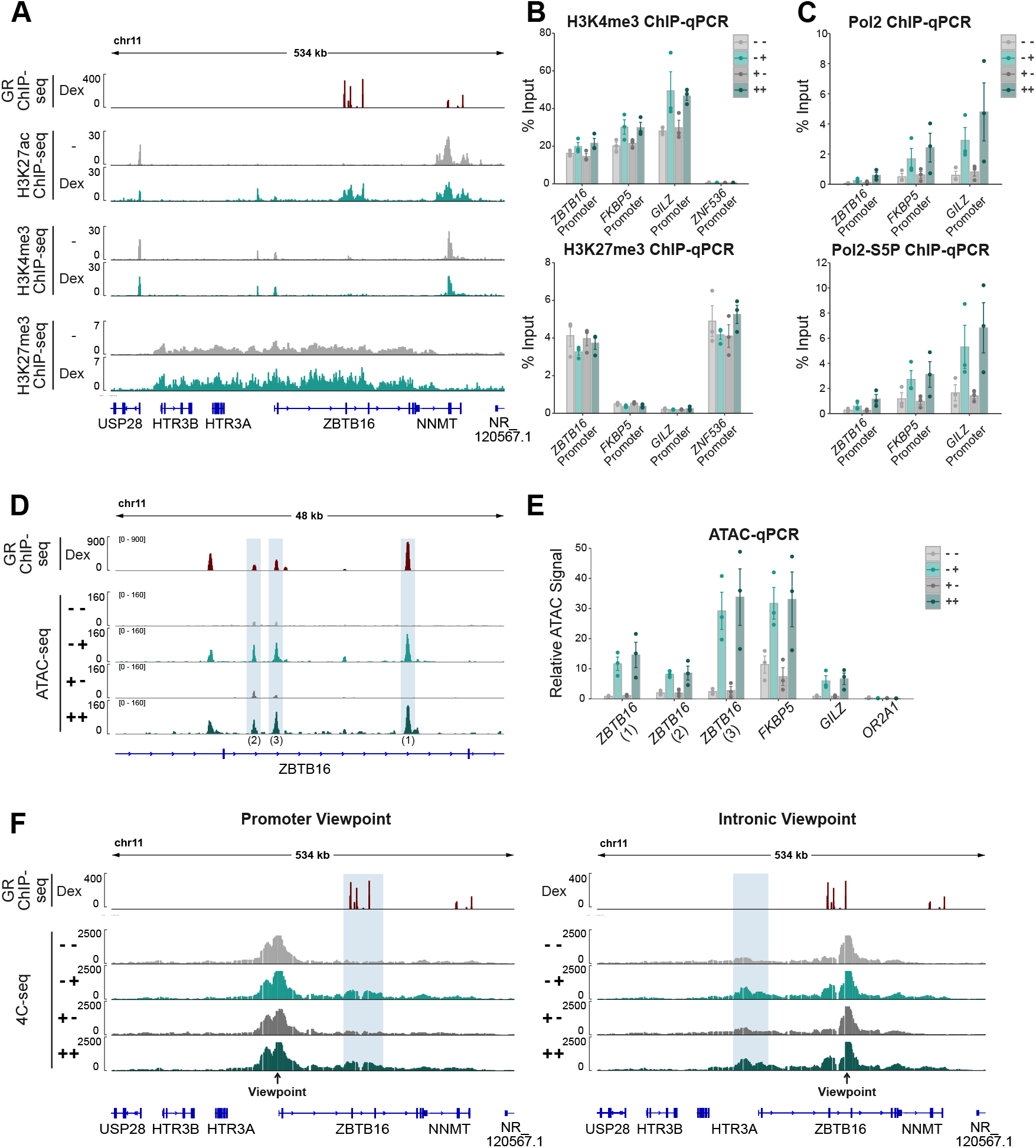
Short-term maintenance of GC-induced chromatin changes at the *ZBTB16* locus. (a) Genome browser visualization of the *ZBTB16* locus in A549 cells showing RPKM-normalized ChIP-seq signal tracks for GR (100 nM Dex, 3h; data from [43]), H3K27ac (+/-100 nM Dex, 4h; data from [44]), H3K4me3 (+/-100 nM Dex, 4h; data from [44]) and H3K27me3 (100 nM Dex or EtOH, 1h; data from [44]). (b) ChIP-qPCR targeting H3K4me3 (top) and H3K27me3 (bottom) at promoters of indicated genes in A549 cells. Cells were treated as described in Fig. 3a. Mean % input ± SEM (n = 3) is shown. (c) Same as for (b) except that ChIP-qPCR targeting RNA PolII (top) and RNA PolII-S5P (bottom) is shown. (d) Genome browser visualization of GR peaks at the *ZBTB16* locus in A549 cells showing RPKM-normalized GR ChIP-seq (100 nM Dex, 3h; data from [43]) and normalized ATAC-seq signal tracks. For the ATAC-seq experiments, the cells were treated as described in Fig. 3a. (e) Highlighted peaks in (d) and other regions as indicated were targeted in ATAC-qPCR quantification. A549 cells were treated as described in Fig. 3a. Mean ATAC signal (normalized to gDNA) ± SEM (n = 3) is shown. (f) Genome browser visualization of the region around the *ZBTB16* locus in A549 cells showing normalized 4C-seq signal tracks with the *ZBTB16* promoter as a viewpoint (left) and an intronic GR peak as a viewpoint (right). Cells were treated as described in Fig. 3a. One representative replicate of two biological replicates is shown. Regions with increased contact frequencies upon GC treatment are highlighted with blue shading.

Taken together, these results suggest that none of the chromatin changes we assayed persist after hormone washout and thus likely do not play a key role in driving transcriptional memory in *ZBTB16* priming.

### Priming increases *ZBTB16* output by increasing the fraction cells responding to hormone treatment and augmented activation by individual cells

A more robust transcriptional response by a population of cells upon a second hormone exposure can be the consequence of a larger fraction of cells responding (Fig. 6a). Moreover, increased transcript levels can be achieved when individual cells respond more robustly when primed (Fig. 6a). To assay *ZBTB16* mRNA expression at a single-cell level, we performed RNA FISH in A549 cells comparing primed and naïve cells. Using probes targeting the coding sequences of *ZBTB16*, we could detect individual transcripts (orange spots in the cytoplasm) as well as the sites of transcription (larger foci in the nucleus, see materials and methods), which reflect the number of actively transcribing alleles (Fig. 6b). Consistent with our RNA-seq and qPCR data, *ZBTB16* levels are barely detectable for untreated cells (Fig. 3e, Fig. 4a, Fig. 6b,c). This changes when cells are treated with hormone, however, increased transcript levels and transcription foci are only detectable for a minor subset of cells (Fig. 6b-d). Conversely, hormone washout resulted in a complete loss of both transcription foci and cells with higher transcript levels (Fig. 6b-d). The response to hormone of primed cells is changed in two ways when compared to naïve cells. First, a larger fraction of cells shows hormone-induced increases in *ZBTB16* transcript levels and transcriptional foci (Fig. 6c,d). Second, a comparison of the distribution of transcripts per cell shows that a subset of the primed cells expresses *ZBTB16* at levels that are higher than for naïve hormone-treated cells (Fig. 6c). These effects were even more pronounced when cells were exposed to GCs a third time (Fig 6c,d). To compare our findings for the *ZBTB16* gene with a GR-target gene that does not change its transcriptional response upon priming, we analyzed the *FKBP5* gene. Compared to *ZBTB16, FKBP5* is expressed at higher basal levels and accordingly its transcripts are detected for the majority of cells regardless of whether cells were hormone-treated or not (Fig. 6e,f). Upon hormone treatment, both the number of transcripts per cell and the number of transcriptional foci increases. However, in contrast to *ZBTB16* the majority of cells respond to hormone treatment by having more transcripts per cell and more detectable transcriptional foci (Fig. 6f,g). Moreover, consistent with a lack of priming, the distribution of transcriptional foci and transcripts per cells for *FKBP5* does not show a noticeable change when comparing naïve and primed cells.

**Figure 6.**
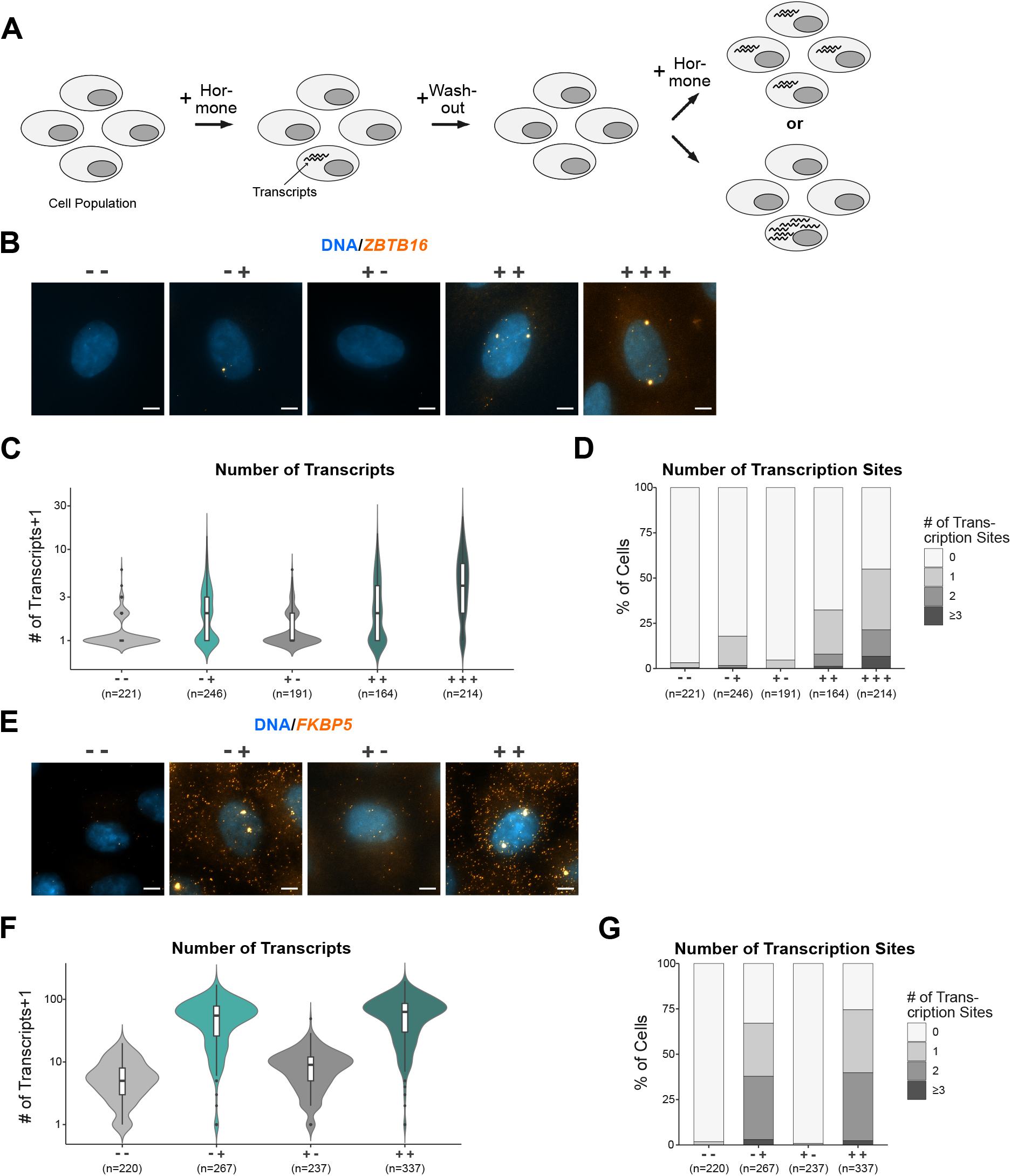
Single-cell analysis comparing the activation of the *ZBTB16* gene between ‘primed’ and ‘naïve’ cells. (a) Cartoon depicting how a more robust response can be driven by both more cells responding and by individual cells that respond more robustly. (b) Representative image of an RNA FISH experiment targeting *ZBTB16* mRNA (orange) in A549 cells. Nuclei were counterstained with DAPI (blue). Cells were hormone treated prior to fixation as described in Fig. 3a. Scale bar 5 μm. (c) Violin plots with box plots inside showing the number of *ZBTB16* transcripts+1 per cell as detected by RNA FISH for treatments as in (b). Results derived from three biological replicates are shown. (d) Stacked bar graphs showing the percentage of cells with 0, 1, 2 or ≥3 visible transcription sites of *ZBTB16* per treatment in (b) as detected by RNA FISH. Results from three biological replicates are shown. (e) Same as (c), except that *FKBP5* mRNA was targeted. (f) Same as (c), except that *FKBP5* mRNA was targeted. (g) Same as (d), except that *FKBP5* mRNA was targeted.

Together, we find that priming changes the number of cells responding and the robustness of the response of individual cells that are specific for the *ZBTB16* gene which together explain the more robust response observed for this gene.

## Discussion

GCs are released by the adrenal cortex in response to various types of stress. For an effective response to acute stress, cells need to react fast but should also reverse their response when the stressor is no longer present. However, when stressors are encountered repeatedly, habituation and an altered, *e*.*g*. blunted, response might be crucial to ascertain an organisms well-being [67]. Physiologically, GCs are released in a circadian and ultradian manner [13,14]. When stress is chronic, this results in a prolonged exposure to high hormone levels and is associated with severe pathological outcomes [10,68]. Similarly, when GCs are used therapeutically, high doses and repeated long-term exposure causes severe side effects and can result in emerging GC resistance [69,70]. Motivated by previous studies that indicated that GCs induce long-term chromatin changes, we explored genome-wide changes in chromatin accessibility and assayed if these changes persist after hormone washout. In addition, we investigated if prior exposure to GCs influences the transcriptional response to the subsequent exposure of the same stimulus. In contrast to previous studies [33,38], we did not find convincing evidence that changes in chromatin accessibility persist after hormone washout as both GC-induced increases and decreases in chromatin accessibility universally reversed to their pre-hormone exposure levels (Fig. 2b). One difference to the prior study that described persisting GR-induced hypersensitivity for more than 9 days is that they studied memory in another cell type (mouse L cell fibroblast) [38]. Another difference is that we studied genome-wide changes, whereas they studied a single exogenous stably integrated MMTV sequence. Hence, it is possible that their results do not represent a phenomenon which is commonly observed at endogenous mammalian loci. Thus, even though we did not find convincing persistence of changes in chromatin accessibility in either one of two cell lines tested, we cannot rule-out that cell type-specific mechanisms facilitate sustained accessibility. A more recent study reported sustained increases in chromatin accessibility in mouse mammary adenocarcinoma cells 40 minutes after hormone washout [33]. The reversal after the 24-hour washout in our study indicates that GC-induced changes are universally short-lived and do not persist beyond one cell cycle in A549 and U2OS cells. We conclude that maintained chromatin openness as a result of GR binding is not a general trait of GR activation and rather appears to represent a cell type- and locus-specific phenomenon.

Mirroring what we say in terms of chromatin accessibility, transcriptional responses also seem universally reversable with no indication of priming-related changes in the transcriptional response to a repeated exposure to GC for any gene with the exception of *ZBTB16*. Although several changes in the chromatin state occurred at the *ZBTB16* locus, none of these changes persisted after hormone washout arguing against a role in transcriptional memory at this locus (Fig. 5). Similarly, the increased long-range contact frequency between the *ZBTB16* promoter region and a GR-occupied enhancer does not persist after washout (Fig. 5e). Notably, our RNA FISH data showed that *ZBTB16* is only transcribed in a subset of cells, hence, it is possible that persistent epigenetic changes occurring at the *ZBTB16* locus also only occur in a small subset of cells and could thus be masked by bulk methods such as ChIP-seq or ATAC-seq. Another mechanism underlying the priming of the *ZBTB16* gene could be a persistent global decompaction of the chromatin as was shown for the *FKBP5* locus upon GR activation [35]. Likewise, sustained chromosomal rearrangements, which we may not capture by 4C-seq, could occur at the *ZBTB16* locus and affect the transcriptional response to a subsequent GC exposure. Furthermore, prolonged exposure to GCs (several days) can induce stable DNA demethylation as was shown for the tyrosine aminotransferase (*Tat*) gene [71]. The demethylation persisted for weeks after washout and after the priming, activation of the *Tat* gene was both faster and more robust when cells were exposed to GCs again [71]. Interestingly, long-term (2 weeks) exposure to GCs in trabecular meshwork cells induces demethylation of the *ZBTB16* locus raising the possibility that it may be involved in priming of the *ZBTB16* gene [72]. However, it should be noted that our treatment time (4 hours) is much shorter. Finally, enhanced *ZBTB16* activation upon a second hormone exposure might be the result of a changed protein composition in the cytoplasm following the first hormone treatment. In this scenario, increased levels of a cofactor produced in response to the first GC treatment would still be present at higher levels and facilitate a more robust activation of *ZBTB16* upon a subsequent hormone exposure. Although several studies have reported gene-specific cofactor requirements [73], the fact that we only observe priming for the *ZBTB16* gene would make this an extreme case where only a single gene is affected by changes in cofactor levels.

ZBTB16 is a transcription factor that belongs to POZ and Krüppel family of transcriptional repressors [74]. It plays a role in various processes including limb development, stem cell self-renewal and innate immune responses [74,75]. Interestingly, both GR and ZBTB16 are linked to metabolic syndrome including changes in insulin sensitivity [76], raising the possibility that the severe metabolic side-effects experienced by patients undergoing long-term GC treatment could involve mis-regulation of *ZBTB16* as a result of repeatedly prolonged GR activation. *ZBTB16* might also play a role in emerging GC-resistance given that increased levels negatively regulate GR transcriptional activity [77]. Specifically, GC-induced apoptosis and gene regulation are blunted upon *ZBTB16* overexpression, whereas GC sensitivity increases when *ZBTB16* is knocked down [78].

In summary, we report that a relatively short-term GR activation results in universally reversable changes in chromatin accessibility as measured by ATAC-seq. This holds true for sites with increased chromatin accessibility but also for a large number of sites we identified that become less accessible upon GC treatment. In contrast to opening sites, closing sites are typically not GR-occupied yet are enriched near repressed genes in line with other studies [23,24,61,64], suggesting that transcriptional repression by GR in general does not require nearby GR binding. Given the circadian release of GCs and their role in responding to stress it makes physiological sense that changes in chromatin accessibility and transcriptional responses are reversible under normal circumstances. However, this might be different when cells are exposed to GCs for extended periods of time, as occurs when GCs are used therapeutically. Despite the universal reversibility of GC-induced chromatin accessibility, we found a single gene that was primed by prior hormone exposure. Interestingly, this gene was only activated in a fraction of cells which is consistent with a recent single-cell RNA-seq study showing that many target genes are only regulated in subset of cells upon GC exposure [79]. Potentially explaining this cell-to-cell variability, single-cell studies show that chromatin states are heterogeneous among populations of cells [80]. Our single-cell studies show that priming resulted in a larger fraction of cells activating *ZBTB16* upon repeated GC treatment and also in a fraction of cells that responded more robustly. Although our bulk studies did not uncover lasting chromatin changes that could explain priming of the *ZBTB16* gene, we envision that single-cell profiling of the chromatin landscape upon repeated exposure to GCs could help further unravel mechanisms that allow individual cells to ‘remember’ a previous hormone exposure and change their response when the signal is encountered again.

## Acknowledgements

We thank Edda Einfeldt and Beatrix Fauler for excellent technical support.

## Author contributions

M.B. performed the experiments and analyzed the data. R.B. and M.B. established protocols for the imaging and automated quantification of the RNA FISH experiments. S.H.M. supervised the study. M.B. and S.H.M. wrote the manuscript.

---

**Figure S1.**
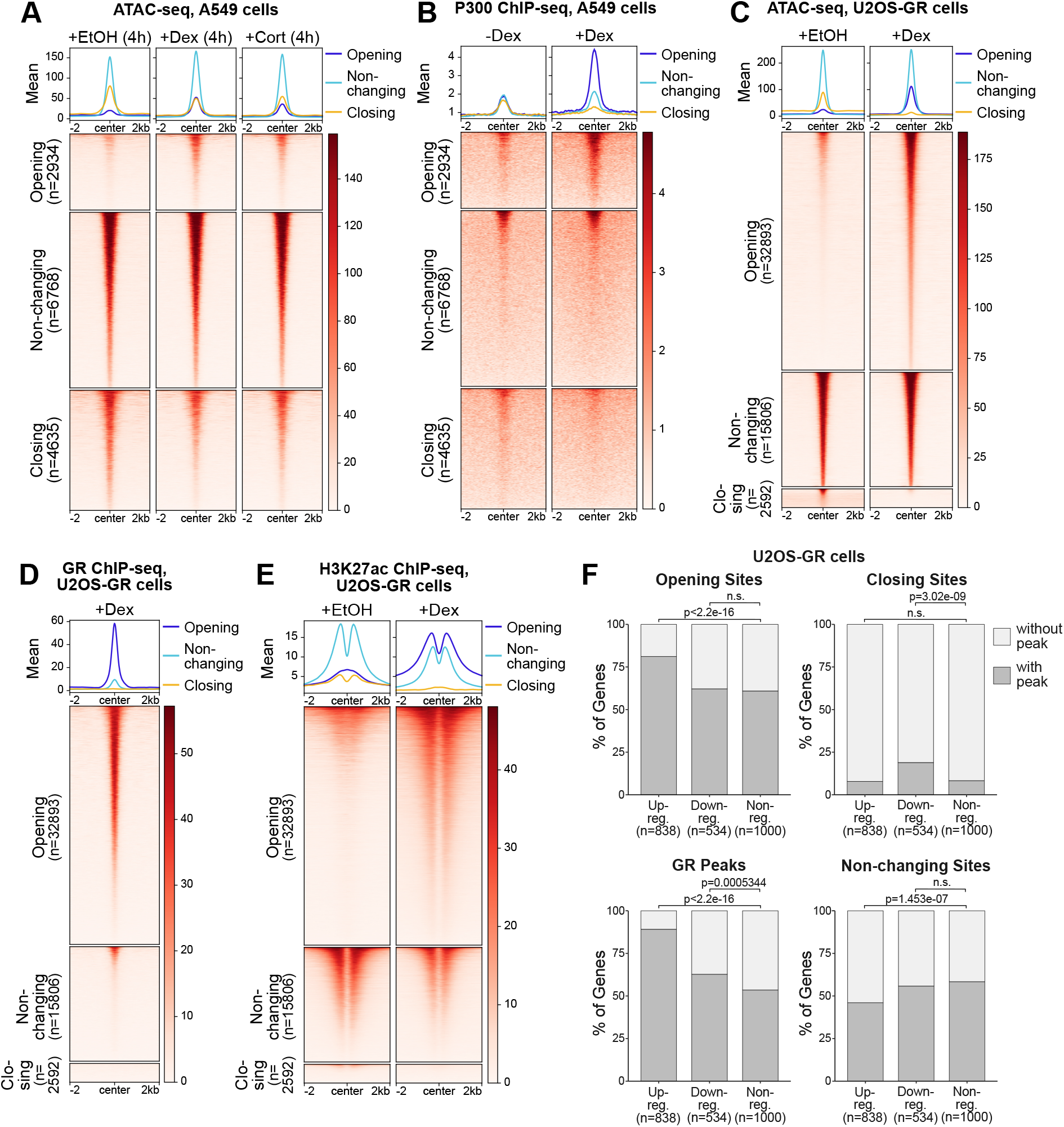
Genome-wide changes in chromatin accessibility in A549 - and U2OS-GR cells upon GR activation. (a) Heatmap visualization and mean signal plot of normalized ATAC-seq read coverage in A549 cells at the same sites as shown in in Fig. 1a (+/-2 kb around center). Cells were treated with EtOH (4h), Dex (100 nM, 4h) or Cort (100 nM, 4h). (b) Same as for (a) except that RPKM-normalized p300 ChIP-seq read coverage (+/-100 nM Dex, 4h) in A549 cells is shown (data from [44]). (c) Heatmap visualization and mean signal plot of normalized ATAC-seq read coverage in U2OS-GR cells at sites of increasing (‘opening’), non-changing (‘non-changing’) and decreasing (‘closing’) chromatin accessibility upon hormone treatment (+/-2 kb around center). Cells were treated with EtOH (4h) or Dex (100 nM, 4h). (d) Same as for (c) except that RPKM-normalized GR ChIP-seq read coverage (1 μM Dex, 1.5h) in U2OS-GR cells is shown (data from [45]). (e) Same as for (c) except that RPKM-normalized H3K27ac ChIP-seq read coverage (1 μM Dex or EtOH, 1.5h) in U2OS-GR cells is shown (data from [46]). (f) Stacked bar graphs showing the percentage of genes in U2OS-GR cells of each category (upregulated, downregulated and nonregulated) that have at least one peak for each type as indicated (opening, closing and non-changing sites and GR peaks) within +/-50 kb around the TSS. P-values were calculated using a Fisher’s exact test. n.s. not significant.

**Figure S2.**
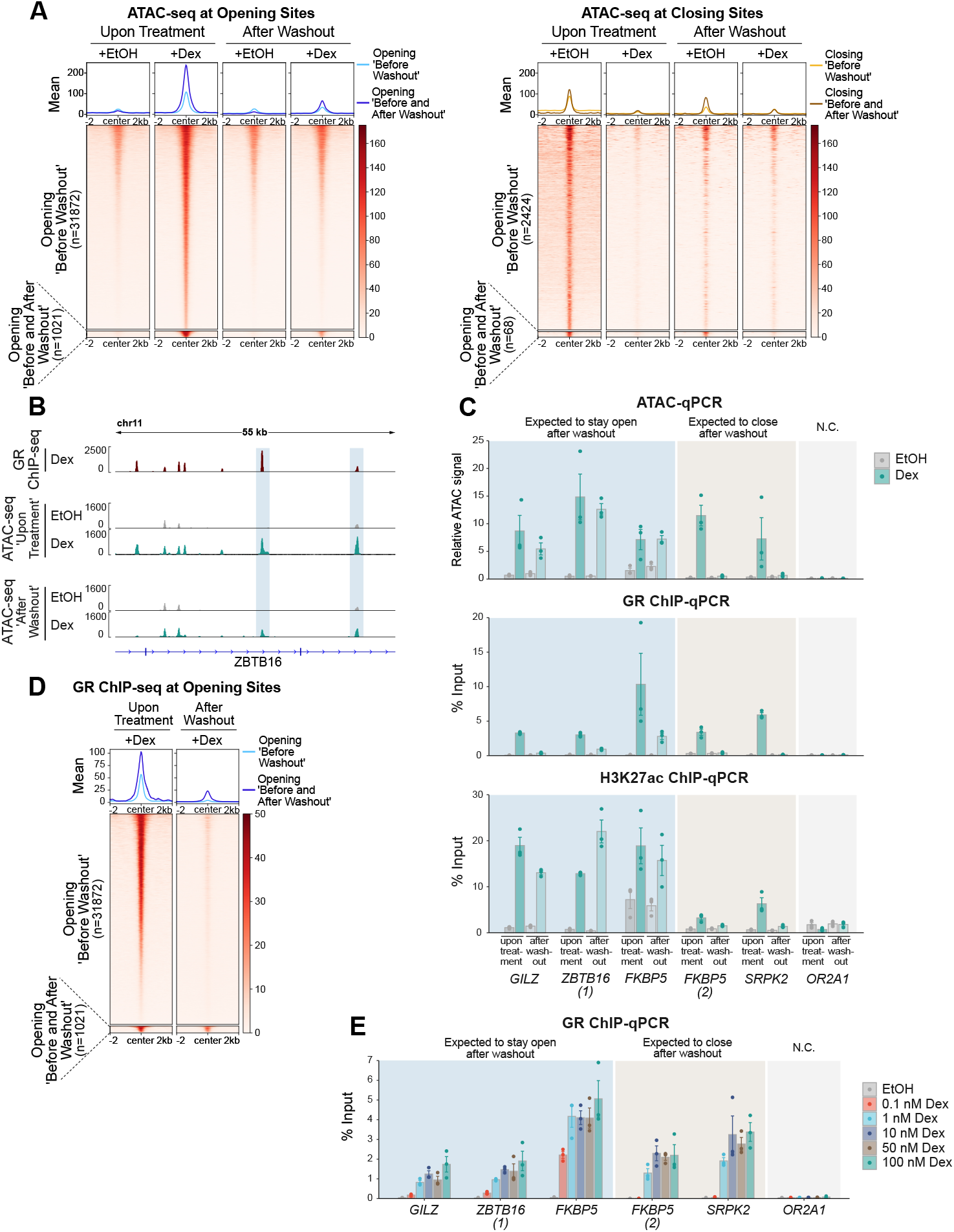
Analysis of GR-induced changes in chromatin accessibility following hormone washout in U2OS-GR cells. (a) Heatmap visualization and mean signal plot of normalized ATAC-seq read coverage in U2OS-GR cells at opening sites (left) and closing sites (right) upon hormone treatment (+/-2 kb around center). Cells were treated similarly as described in Fig. 2a, except that cells were treated (1) with Dex (100 nM) or EtOH for 4h (‘upon treatment’) and (2) with Dex (100 nM) or EtOH for 4h, followed by washes and subsequent culturing in hormone-free medium for 24h (‘after washout’). The regions are divided into sites which show increased/decreased accessibility upon hormone treatment and regions which show increased/decreased accessibility that persists after washout. (b) Genome browser visualization of the *ZBTB16* locus in U2OS-GR cells showing GR ChIP-seq (1 μM Dex, 1.5h; RPKM-normalized; data from [45]) and ATAC-seq (normalized) signal tracks. For ATAC-seq, cells were treated as described in (a). Sites of increased accessibility after hormone washout are highlighted with blue shading. (c) ATAC-qPCR (top), GR ChIP-qPCR (middle) and H3K27ac ChIP-qPCR (bottom) of sites opening upon hormone treatment near indicated genes in U2OS-GR cells. Cells were treated as described in (a). Regions which are expected (based on the ATAC-seq data) to close or remain accessible following hormone washout are indicated. Mean ATAC signal (normalized to gDNA) or mean % input ± SEM (n = 3) is shown. (d) Heatmap visualization and mean signal plot of RPKM-normalized GR ChIP-seq read coverage in U2OS-GR cells at regions of increasing chromatin accessibility (same as ‘opening sites’ shown in (a)) (+/-2 kb around center). For the GR ChIP-seq ‘upon treatment’, cells were treated with Dex (1 μM, 1.5h; data from [45]). For the GR ChIP-seq ‘after washout’, cells were treated with Dex (100 nM, 4h), followed by washes and culturing in hormone-free medium for 24h. (e) GR ChIP-qPCR at same regions as in (c). Cells were treated for 4h with EtOH or Dex at different concentrations as indicated (0.1 nM, 1 nM, 10 nM, 50 nM and 100 nM). Mean % input ± SEM (n = 3) is shown.

**Figure S3.**
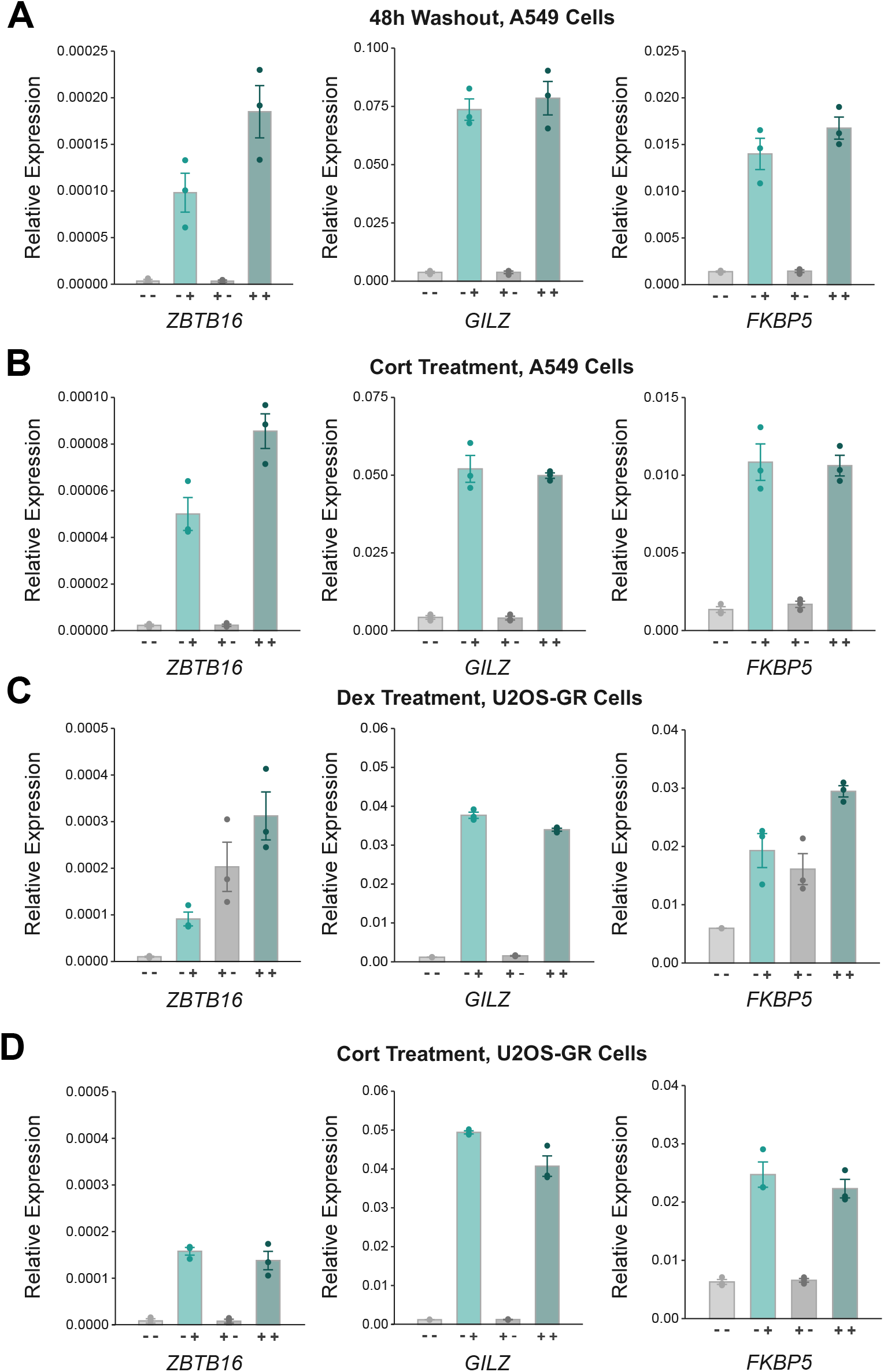
Priming of *ZBTB16* is cell type-specific. (a) RT-qPCR results of *ZBTB16* in A549 cells. Cells were treated as described in Fig. 3a, except that the washout period was 48h instead of 24h. Mean transcript levels relative to *RPL19* ± SEM (n = 3) are shown. (b) RT-qPCR results of *ZBTB16* in A549 cells. Cells were treated as described in Fig. 3a, except that 100 nM Cort was used instead of Dex. Mean transcript levels relative to *RPL19* ± SEM (n = 3) are shown. (c) RT-qPCR results of *ZBTB16* in U2OS-GR cells. Cells were treated as described in Fig. 3a. Mean transcript levels relative to *RPL19* ± SEM (n = 3) are shown. (d) RT-qPCR results of *ZBTB16* in U2OS-GR cells. Cells were treated as described in Fig. 3a, except that 100 nM Cort was used to treat cells instead of Dex. Mean transcript levels relative to *RPL19* ± SEM (n = 3) are shown.

